# An intrinsic alkalization circuit turns on *mntP*-riboswitch under manganese stress in *Escherichia coli*

**DOI:** 10.1101/2022.08.31.505976

**Authors:** Arunima Kalita, Rajesh Kumar Mishra, Vineet Kumar, Amit Arora, Dipak Dutta

## Abstract

The trace metal manganese in excess affects iron-sulfur cluster and heme-protein biogenesis eliciting cellular toxicity. The manganese efflux protein MntP is crucial to evading manganese toxicity in bacteria. Recently, two Mn-sensing riboswitches upstream of *mntP* and *alx* in *Escherichia coli* have been reported to mediate the upregulation of their expression under manganese shock. As the *alx*-riboswitch is also responsive to alkaline shock administered externally, it is intriguing whether *mntP*-riboswitch is also responsive to alkaline stress. Furthermore, how both manganese and alkaline pH simultaneously regulate these two riboswitches under physiological conditions is a puzzle. Using multiple approaches, we show that manganese shock activated glutamine synthetase (GlnA) and glutaminases (GlsA and GlsB) to spike ammonia production in *E. coli*. The elevated ammonia intrinsically alkalizes the cytoplasm. We establish that this alkalization under manganese stress is crucial for attaining the highest degree of riboswitch activation. Additional studies showed that alkaline pH promotes a 17 to 22-fold tighter interaction between manganese and the *mntP*-riboswitch element. Our study uncovers a physiological linkage between manganese efflux and pH homeostasis that mediates enhanced manganese tolerance.

**Significance statement:** Riboswitch RNAs are cis-acting elements that can adopt alternative conformations in the presence or absence of a specific ligand(s) to modulate transcription termination or translation initiation processes. In the present work, we show that how manganese and alkaline pH both are necessary for maximal *mntP*-riboswitch activation to mitigate the manganese toxicity. This study bridges the gap between earlier studies that separately emphasize the importance of alkaline pH and manganese in activating the riboswitches belonging to the *yybP-ykoY*-family. This study also ascribes a physiological relevance as to how manganese can rewire cellular physiology to render cytoplasmic pH alkaline for its homeostasis.

## Introduction

Manganese is a crucial metal that determines the pathogenic potential of bacteria (1-3). This metal has been shown to sustain the functions of a small subset of proteins, such as superoxide dismutase and ribonucleotide reductase, under iron starvation and oxidative stress (4-6). Conversely, iron starvation and oxidative stress are also induced in *E. coli* in a laboratory setting by the presence of excess manganese in the medium (7-9). Excess manganese at a toxic level impairs the biogenesis of iron-sulfur cluster and heme-containing proteins in *E. coli*, thereby compromising the function of proteins, involved in the electron transport chain, affecting energy metabolism (8-10).

A manganese-dependent transcription regulator (MntR) downregulates the manganese importer (*mntH*) and upregulates the manganese exporter (*mntP*) to alleviate manganese toxicity in *E. coli* (7). Notably, MntP-mediated efflux of manganese alone efficiently controls the detrimental effects of manganese overload (7-9). As a result, Δ*mntP* strain is more sensitive to manganese than WT *E. coli* cells (7). Interestingly, a recent study has demonstrated that a 5’-untranslated region (5’-UTR) of *mntP* transcript forms a manganese-dependent riboswitch to turn on translation initiation of *mntP*, causing successful manganese efflux (11).

Riboswitch RNAs can adopt at least two alternative conformations depending on specific ligand(s) availability to modulate transcription termination or translation initiation processes (12). The *mntP* riboswitch and another riboswitch at 5’-UTR of *alx* gene in *E. coli* belong to the *yybP-ykoY* riboswitch family that has been shown to be regulated by manganese (11, 13-15). However, the preceding reports suggest that the *alx*-riboswitch is also responsive to alkaline shock administered externally (11, 16, 17). In this context, it is intriguing whether *mntP* riboswitch also responds to alkaline stress.

We previously demonstrated that manganese stress inhibits the activity of glutamate synthase (GOGAT), an iron-dependent enzyme (9). Together with this information, our present work reveals that manganese shock intrinsically spikes cellular ammonia level by altering the activities of the glutamate-glutamine cycle enzymes, viz. GOGAT, glutamine synthetase (GlnA), and glutaminases (GlsA and GlsB) enzymes. This ammonia production elevates cellular pH favoring the activation of the *mntP* riboswitch.

## Results

### Glutamine synthetase function is critical for manganese homeostasis

Glutamine synthetase (GlnA) utilizes Mn^2+^ as a cofactor (18). Our earlier finding has shown that the level of GlnA is increased in the manganese-fed *E. coli* Δ*mntP* strain (9). We observed that 8 mM and 1 mM manganese caused similar growth inhibition in WT, and Δ*mntP* strains, respectively (Figure S2A). This observation suggests that the degree of manganese toxicity in 8 mM manganese-treated WT strain is similar to the 1 mM manganese-treated Δ*mntP* cells. Next, we performed western blotting to show that the elevated GlnA levels were comparable in the WT and Δ*mntP* strains treated with 8 mM and 1 mM manganese, respectively (Figure 1A).. Further growth analysis reveals that 0.5 mM manganese was severely toxic to the Δ*mntP*Δ*glnA* strain in comparison to the Δ*mntP* or Δ*glnA* single mutants (Figure 1B; Figure S1). This toxicity is rescued by the presence of the plasmid pGlnA that expressing functional *glnA* (Figure S2B). Furthermore, we observed that the viability of the Δ*mntP*Δ*glnA* double mutant was 3-times lesser than the Δ*mntP* strain under a lethal higher dose of manganese (10 mM), that was alleviated by expression of *glnA* from the pGlnA plasmid (Figure S2C). These observations suggest that GlnA function is crucial for the Δ*mntP* strain under manganese shock.

**Figure 1.**
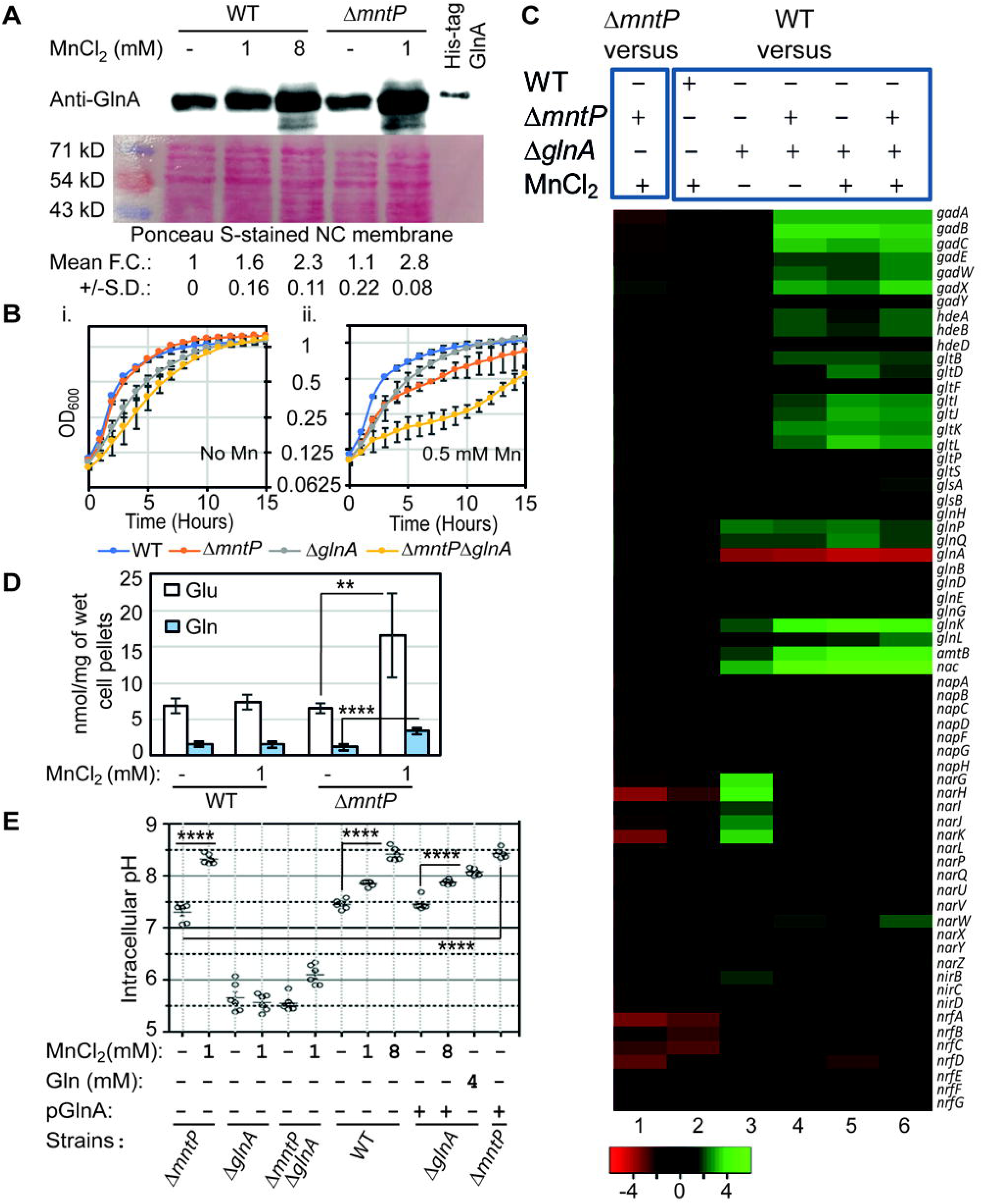
Manganese-mediated GlnA activation triggers intracellular pH elevation. **A**. Western blot shows upregulated GlnA in the manganese-fed WT and Δ*mntP* strains. The corresponding ponceau-stained nitrocellulose membrane was represented to show equal loading of the cellular proteins. The fold change (FC) values are the mean ± S.D. from three independent blots. **B**. Growth curves show that Δ*mntP*Δ*glnA* double mutant is extremely sensitive to 0.5 mM manganese than the single mutants. **C**. The microarray heat-map representing up- and downregulated genes. **D**. Intracellular levels of glutamate and glutamine in the presence or absence of manganese stress. **P <0.01, ****P <0.0001. **E**. A GFP reporter system (GFPmut3*) was used to determine the intracellular pH. The intracellular pH in the presence or absence of manganese, glutamine or pGlnA plasmid were estimated. ****P < 0.0001. The graphs were plotted from mean ± S.D. from six different experiments.

To understand how *glnA* contributes to overcoming manganese toxicity, we performed microarray experiments using WT, Δ*glnA* and Δ*mntP*Δ*glnA* mutants grown in the presence or absence of 1 mM manganese. The altered gene expressions in respect to the WT strain were identified and compared with our previously reported altered gene expression profile of the Δ*mntP* cells treated with manganese (Figure 1C) (9). While most of the genes for glutamate/aspartate or glutamine transporters (*gltIJKL, gltP*, and *glnHPQ*) were upregulated, the essential genes involved in glutamate-dependent acid resistance (*e*.*g*., *gadABC*) and nitrate/nitrite reductase pathways (*nap, nar, nir* and *nrf* genes) were repressed in the manganese-fed Δ*mntP* strain (Figure 1C). A modest level of repression of the genes in the nitrate/nitrite reductase pathways in the manganese-fed WT strain was also observed, suggestive of a mild effect of 1 mM manganese in the WT cells (Figure 1C). These observations indicate that the intracellular pH homeostasis and nitrate/nitrite metabolism could be altered under manganese toxicity.

On the other hand, expressions of acid resistance, nitrate/nitrite metabolism, and ammonia transport (*amtB*) genes were upregulated in Δ*glnA* strain (Figure 1C). The genes that confer acid resistance, glutamate-glutamine cycling, and ammonia transport were further strongly activated in Δ*mntP*Δ*glnA*, and manganese-fed Δ*glnA* and Δ*mntP*Δ*glnA* strains (Figure 1C). Interestingly, the expression of the genes in the nitrate/nitrite reductase pathways were sequentially decreased in the following order: Δ*glnA*> manganese-fed Δ*glnA*> Δ*mntP*Δ*glnA*> manganese-fed Δ*mntP*Δ*glnA* (Figure 1C). These observations indicate that GlnA possibly alters nitrogen metabolism and pH homeostasis to alleviate manganese shock.

### GlnA-mediated glutamine synthesis is crucial for the increased cellular pH under manganese toxicity

The glutamate-glutamine cycle generates glutamate and glutamine, which produce all other nitrogenous metabolites (19). Glutamate and glutamine play pivotal roles in acid-resistant systems (20-22). We noted that the cellular pool of these two amino acids increased up to 3-fold in the manganese-treated Δ*mntP* strain (Figure 1D). Since manganese stress blocks heme biogenesis (8), where glutamate acts as a precursor, we can argue that fluxes of glutamate to heme will be inhibited, causing increased glutamate level under manganese stress. From this observation, we speculate that the activated GlnA (Figure 1, A and B) increased the cellular glutamine pool in the manganese-fed Δ*mntP* strain.

We have previously noticed that the cellular pH in the manganese-fed Δ*mntP* strain is elevated (9). In this study, we accurately determined intracellular pH in the unfed and manganese-fed strains of *E. coli* using a pH-sensitive GFPmut3*-bearing plasmid (23, 24). Δ*mntP* mutant exhibited a cellular pH of 7.3 ± 0.18, and it was dramatically elevated to about pH 8.3 ± 0.09 under 1 mM manganese treatment (Figure 1E). However, the cellular pH was declined to 5.7 ± 0.27 and 5.5 ± 0.15 in the Δ*glnA* and Δ*mntP*Δ*glnA* strains, respectively (Figure 1E). Further exposure to 1 mM manganese failed to elevate the cellular pH in Δ*glnA* strain (Figure 1E). The intracellular pH of Δ*mntP*Δ*glnA* strain modestly increased to 6.1 ± 0.2 upon 1 mM manganese exposure (Figure 1E). Intracellular pH of the WT strain was elevated from 7.4 ± 0.08 to 7.8 ± 0.05 and 8.4± 0.12 under 1 mM and 8 mM manganese exposure, respectively (Figure 1E).

The Δ*glnA* strain was complemented with a plasmid bearing *glnA* (pGlnA) to show that the cellular pH was elevated to 7.4 ± 0.11, which was further raised to 7.9 ± 0.04 upon exposure to 1 mM manganese, a value similar to the pH of 1 mM manganese-fed WT strain (Figure 1E). Interestingly, expression of *glnA* from pGlnA alone could elevate pH to 8.4± 0.09 in the Δ*mntP* strain without any manganese shock (Figure 1E). We also supplemented 4 mM glutamine to show that it elevated the intracellular pH to 8.1 ± 0.06 in the Δ*glnA* strain (Figure 1E). These data additionally support the observation that the GlnA-mediated glutamine production plays a crucial role in elevating intracellular pH under manganese stress.

### Glutaminases catalyze ammonia production to raise cellular pH under manganese stress

From the involvement of GlnA and the upregulation of *amtB* in Δ*glnA* and Δ*mntP*Δ*glnA* strains (Figure 1C), we speculate that a likely elevation of ammonia levels might be a factor contribution to the surge in pH under manganese stress. Indeed, cellular ammonia level was found to be increased in the Δ*mntP* strain (Figure 2A). Manganese supplementation further elevated ammonia in Δ*mntP* strain (Figure 2A). The ammonia levels in Δ*glnA* and Δ*mntP*Δ*glnA* strains remained at basal WT level (Figure 2A). Interestingly, 1 mM manganese treatment raised the ammonia level in the Δ*mntP*Δ*glnA* strain (Figure 2A), plausibly due to the sole utilization of cellular glutamine through the GlsA/B enzymes to produce glutamate and ammonia, as manganese treatment inactivates GOGAT to produce glutamate (9). The last observation also explains the modestly elevated pH in the manganese-fed Δ*mntP*Δ*glnA* strain (Figure 2A).

**Figure 2.**
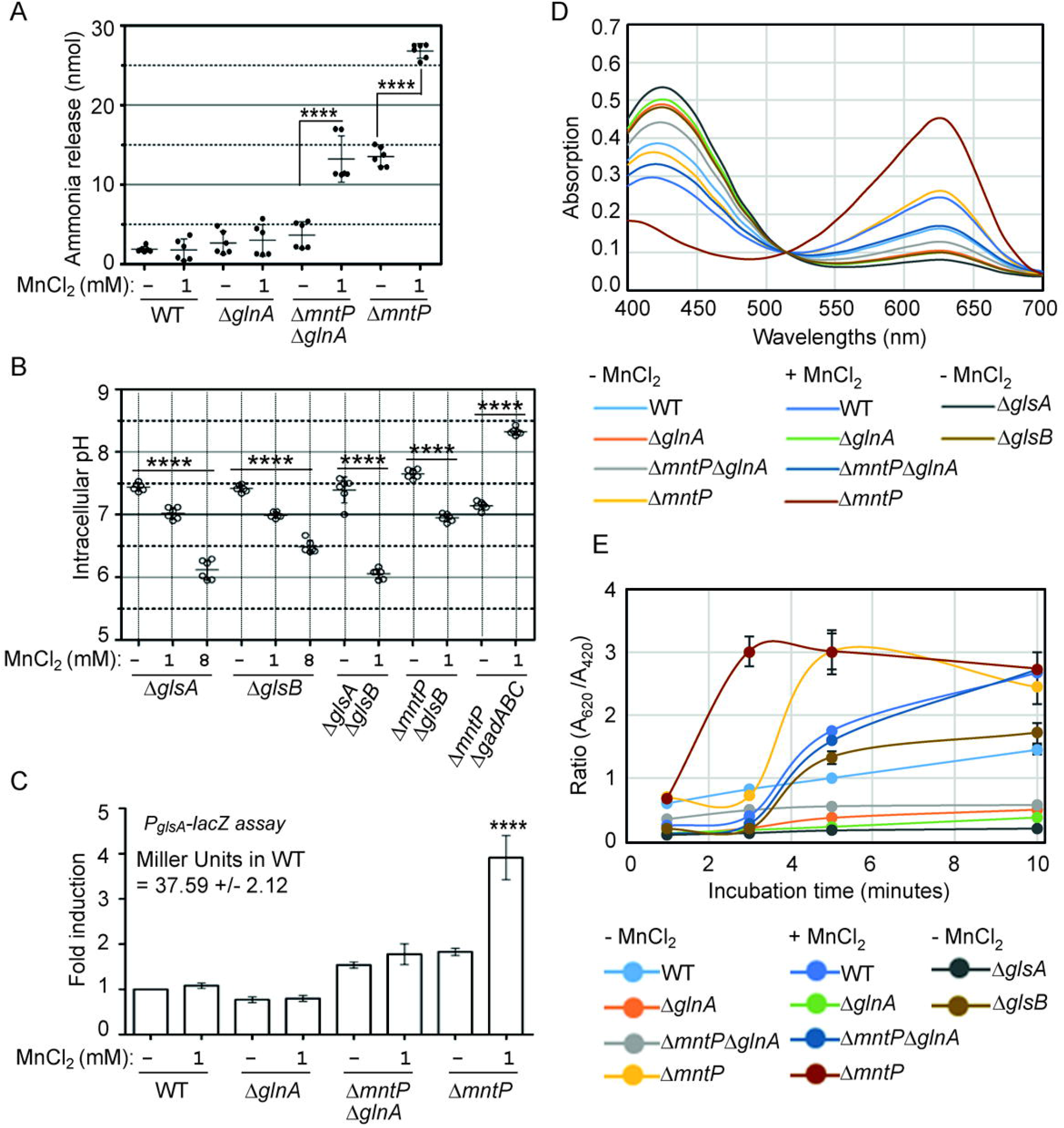
Ammonia liberation by GlsA/B raises intracellular pH. **A**. Cellular ammonia release from the unfed and manganese-fed *E. coli* strains was estimated using ab83360 Ammonia Assay Kit (Abcam) and plotted. ****P < 0.0001. **B**. Intracellular pH was measured and plotted for the *E. coli* strains grown in the presence or absence of manganese using GFPmut3* reporter. ****P <0.0001. **C**. Relative activity of *glsA*-promoter in the different *E. coli* strains grown in the presence or absence of manganese was plotted (using a lacZ reporter). The promoter activity in WT strain was 37.59 ± 2.12 Miller units, which was normalized to 1 for fold-change calculation. ****P <0.0001 against untreated WT control. **D**. Absorption spectrums of bromocresol green pH indicator dye was recorded after allowing the cells to liberate ammonia. Note that the absorption at 620 nm sharply increased for Δ*mntP* strain treated with manganese at 3 minutes time-point (also see Figure S5A for all time-points). **E**. A_620_/A_420_ of bromocresol green dye was obtained and plotted to show the kinetics of ammonia release from different strains grown in the presence or absence of manganese.

The GlsA and GlsB enzymes break down glutamine to produce glutamate, releasing ammonia (21). We assessed that Δ*glnA*Δ*glsA*, Δ*mntP*Δ*glsA*, and Δ*mntP*Δ*glnA*Δ*glsA* strains grow poorly compared to Δ*glnA*, Δ*glsA*, and Δ*mntP* single mutants on LB agar (Figure S3). Besides, the double and triple mutants formed partially lysed cell pellets while growing in LB broth. Δ*glnA*Δ*glsB* and Δ*mntP*Δ*glnA*Δ*glsB* mutants also grew poorly on LB agar (Figure S3), but they were not lysed in LB broth. Δ*mntP*Δ*glsB* strain grew optimally both on LB agar and LB broth. These observations indicate that the GlsA function is more critical than GlsB in Δ*mntP* mutant.

We found that the Δ*glsA* and Δ*glsB*, and Δ*mntP*Δ*glsB* strains exhibited an optimum physiological pH. 1 mM manganese modestly declined cytoplasmic pH to 7.0 ± 0.09 (Figure 2B). 8 mM manganese declined the pH to 6.1 ± 0.16 and 6.5 ± 0.1 in Δ*glsA* and Δ*glsB* mutants, respectively (Figure 2B). On the other hand, 1 mM manganese was enough to drop the intracellular pH to 6.1 ± 0.08 in the Δ*mntP*Δ*glsB* strain. Why manganese stress caused the pH drops in Δ*glsA*, Δ*glsB*, and Δ*mntP*Δ*glsB* mutants is discussed later. These observations indicate that manganese fails to elevate the intracellular pH in the absence of GlsA or GlsB. The glutamate-dependent acid resistant (*gad*) system did not play any role in alkalization in Δ*mntP* mutant, as evident from the observation that Δ*mntP*Δ*gadABC* mutant successfully elevated the pH to 8.3± 0.06 (Figure 2B).

To check whether *glsA* was upregulated under manganese stress, we constructed a *glsA-lacZ* transcriptional fusion (P_*glsA*_-*lacZ*), and integrated it into the genome of the strains under study. Δ*mntP* and Δ*mntP*Δ*glnA* strains exhibited about 1.7-fold increase in β-galactosidase activity compared to WT counterpart (Figure 2C). Upon exposure to 1 mM manganese, Δ*mntP*, but not Δ*mntP*Δ*glnA* strain, exhibited a 3.7-fold increase in β-galactosidase activity (Figure 2C). The *glsA-lacZ* activity remained unaltered in unfed and manganese-fed Δ*glnA* cells (Figure 2C).

We performed a GlsA activity assay using the bromocresol green pH indicator dye. Instead of merely visualizing color changes (Figure S4A), as described (25), we recorded the absorption spectrum of the dye (Figure S4B). A gradual increase in the ratio of the two absorption peaks (A_620_/A_420_) of the dye highly correlated with the increasing amounts of an added base to raise the pH from 3.2 to 5.9 (Figure S4C). Thus, we consider the GlsA-catalyzed ammonia base liberation from *E. coli* strains as a function of A_620_/A_420_.

A steep increase in A_620_ and A_620_/A_420_ values was observed within 3-minutes when the indicator dye was incubated with unfed Δ*mntP* or manganese-fed Δ*mntP* cells (Figure 2D and 2E), indicating a faster ammonia release under this condition. As expected, A_620_/A_420_ ratio suggested that the Δ*glsA* strain did not liberate any ammonia with time (Figure 2E). Similarly, from the A_620_/A_420_ ratio, it appeared that unfed and manganese-fed Δ*glnA* and unfed Δ*mntP*Δ*glnA* strains also did not liberate ammonia, indicating that GlsA function in these strains was compromised (Figure 2, D and E). However, manganese-fed Δ*mntP*Δ*glnA* mutant raised A_620_/A_420_ value to some extent, indicating a moderate GlsA activity in this condition (Figure 2E). This observation is consistent with the finding that intracellular ammonia levels increased in the manganese-fed Δ*mntP*Δ*glnA* strain (Figure 2A). Unfed WT and Δ*glsB* strains increased the A_620_/A_420_ ratios very slowly, while manganese-fed WT strain moderately elevated A_620_/A_420_ values throughout the incubation (Figure 2E). We attempted to correlate ammonia liberation as a function of pH changes. However, unlike alterations in A_620_/A_420_ with the increasing amounts of added bases, the alterations in pH units were negligible (Figure S5), as bromocresol green dye has a pKa value of ∼5.0 (Figure S4).

### Intracellular alkalization dictates manganese-driven riboswitch activation

We determined the cellular manganese concentrations in *E. coli* strains (Figure 3A). 1 mM manganese-fed Δ*mntP* strain retained the highest level of manganese (Figure 3A). WT cells treated with 1 mM exogenous manganese elevated the cellular level of manganese to the extent that was equivalent to the manganese content in the unfed Δ*mntP* strain (Figure 3A). Exogenous 8 mM manganese raised the cellular manganese level to 400 μM in the WT cells (Figure 3A). Since 1 mM and 8 mM manganese conferred similar phenotypes to the Δ*mntP* and WT strains, respectively (Figure 1A; Figure 1E; Figure S2A), we suppose that ∼400 μM would be a threshold level of cellular manganese that confers maximum degree of toxicity in *E. coli*. We estimated that 1 mM manganese-fed Δ*mntP*Δ*glnA* and Δ*mntP*Δ*glsB* strains, and 8 mM manganese-fed Δ*glsA* and Δ*glsB* strains accumulated more than or equal to the threshold level of manganese (Figure 3A).

**Figure 3.**
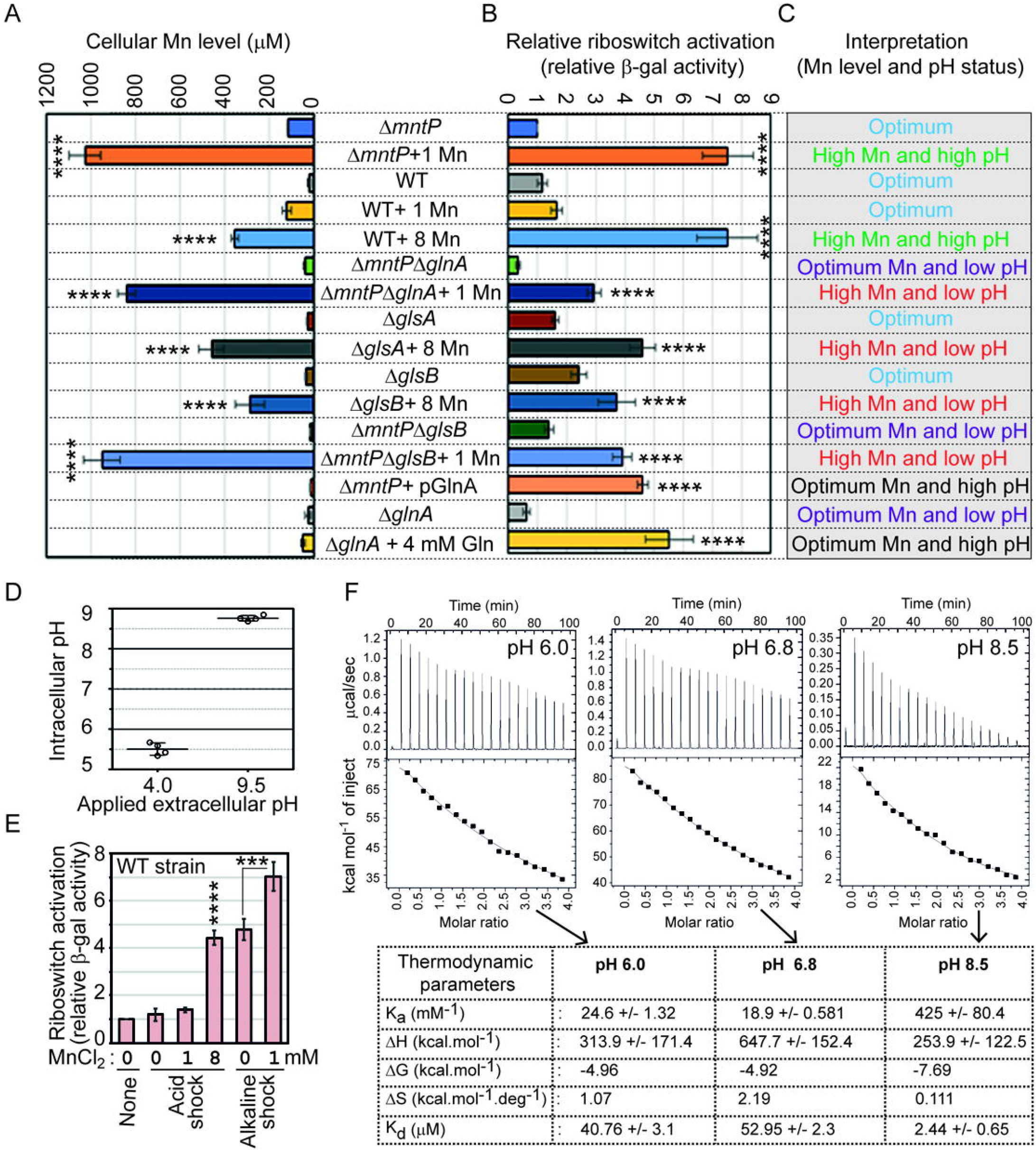
The concerted action of cellular manganese and alkaline pH to activate the *mntP*-riboswitch. **A**. Bar diagram represents the estimated levels of intracellular manganese in the untreated and treated *E. coli* strains (using ICP MS methodology). ****P <0.0001 against untreated Δ*mntP* strain. **B**. Relative riboswitch activity in the different strains grown under specific conditions were plotted from the lacZ reporter (β-gal) assays. The riboswitch activity in Δ*mntP* strain was 16.62 ± 1.62 Miller units, which was normalized to 1 for the fold-change calculation. ****P <0.0001 against untreated Δ*mntP* strain. **C**. Combining the cellular pH (Figure 1E and 2B) and manganese levels (Figure 3A) in a specific strain grown under desired conditions, we classified intracellular environments into five different categories. **D**. Intracellular pH of WT *E. coli* strain was determined under the specific acid and alkaline conditions. **E**. Relative riboswitch activity was estimated in the WT *E. coli* strain under acid and alkaline condition and in the presence or absence of manganese. The riboswitch activity in WT strain was 16.84 ± 1.01 Miller units, which was normalized to 1 for the fold-change calculation. ***P <0.001; ****P <0.0001 against untreated WT strain. **F**. ITC experiments were performed under 3 different pH, viz. 6.0, 6.8, and 8,5, to get binding parameters for riboswitch and manganese interaction.

Recently, a *lacZ-*reporter-construct has been used to demonstrate the riboswitch activity in an *mntP*-deficient PM1205 *E. coli* strain under manganese shock (11). Similarly, we constructed a P_T7A1_ -5′-UTR_*mntP*_ –*lacZ* reporter. Consistent with the observation (11), our reporter-construct indicated a 7.5-fold increase in β-galactosidase activity in 1 mM manganese-fed and 8 mM manganese-fed Δ*mntP* and WT strains, respectively (Figure 3B).

Combining the levels of cellular manganese (Figure 3A) and pH (Figure 1E and Figure 2B) in *E. coli* strains, we categorized intracellular environments as (i) ***Optimum*:** when intracellular manganese was below the threshold level, and pH remained around 7.4 (ii) ***High Mn and high pH*:** when intracellular manganese was equal to or above the threshold level, and pH elevation was above 8.0 (iii) ***High Mn and low pH*:** when intracellular manganese was equal to or above threshold level, but pH was declined to be acidic (iv) ***Optimum Mn and high pH***: when intracellular manganese remained at optimum but pH elevation was above 8.0 (v) ***Optimum Mn and low pH***: when intracellular manganese remained at optimum but pH was declined to be acidic (Figure 3C). As expected, the “high Mn and high pH” category (1 mM and 8 mM manganese-fed Δ*mntP* and WT strains, respectively) showed the highest level of riboswitch activation (Figure 3, B and C). Interestingly, in the “high Mn and low pH” category (represented by 1 mM manganese-fed Δ*mntP*Δ*glnA* and Δ*mntP*Δ*glsB* or 8 mM manganese-fed Δ*glsA and* Δ*glsB* strains), the riboswitch activity remained partial compared to the “*high Mn and high pH*” category (Figure 3, B and C). Similarly, in the “optimum Mn and high pH” category (represented by Δ*mntP* strain harboring pGlnA and Δ*glnA* strain fed with 4 mM glutamine), the riboswitch was activated substantially but not to the highest level (Figure 3, B and C). This data suggests that both alkalization or manganese toxicity can partially activate the riboswitch, but to achieve the highest level of activation, concerted action of manganese and cytoplasmic alkalization was required. To recreate this scenario, we grew WT strain under acidic or alkaline shock (pH 4.0 or 9.5; unbuffered) so that the cellular pH equilibrated to 5.5 or 8.6, respectively (Figure 3D) and subsequently observed the effect of supplemental manganese on the riboswitch activation. Bolstering our finding, the alkaline pH, or the toxic level of manganese at acidic pH could partially activate the riboswitch, but the combination of both activated the riboswitch to the maximal level (Figure 3E). We also performed the riboswitch activity in different pH (buffered) media in the presence or absence of manganese in the Δ*mntP* strain. Consistently, high pH (8.5) alone could activate the riboswitch to 5-fold, which further increases to 9-fold in the presence of manganese (Figure S6A). However, low pH (pH 5.0) and neutral pH (pH 7.0) media could not activate the riboswitch sufficiently in the presence of 1 mM manganese (Figure S6A). These data directly establishes that both alkaline pH and manganese have additive effects in the activation of *mntP*-riboswitch. As a negative control, we used P_T7A1_-lacZ reporter to show that the promoter activity was not influenced by pH variations (Figure S6B).

Similarly, we incorporated *alx*-riboswitch reporter (P_T7A1_ -5′-UTR_*alx*_ –*lacZ* reporter) in theΔ*mntP* strain to show that 1 mM and 2 mM exogenous manganese activated *alx* riboswitch to 3- and 6.5-folds, respectively in unbuffered LB broth, where manganese treatment could intrinsically raise the pH (Figure S7). This manganese-mediated riboswitch activation was dramatically suppressed in buffered LB broth at pH 5.0 (Figure S7). However, the riboswitch activation was modest (3.5-fold under 2 mM manganese stress) in buffered LB broth at pH 7.0 (Figure S7). Remarkably, the *alx*-riboswitch was activated at highest level in the presence manganese in buffered LB broth at pH 8.5 (Figure S7). Interestingly, pH 8.5 alone could activate the riboswitch to 3.5-folds (Figure S7). These data support the previous reports that either alkaline pH or manganese can activate the *alx*-riboswitch (11, 16, 17). Herein, we show that cumulative effect of both alkaline pH and manganese is required for *mntP*-and *alx*-riboswitch activation at highest degrees.

We conducted ITC experiments to address whether the alkaline pH sensitizes the interaction between manganese and the riboswitch element. For this study, we generated the whole 5’-UTR of *mntP* using T7 RNA polymerase-based *in vitro* transcription. The RNA was dissolved in buffers with pH 6.0, 6.8, and 8.5 to equilibrate. 4 μM RNA was titrated by increasing concentrations of MnCl_2_ at 25°C. The ITC profiles exhibited an entropy-driven reaction at pH 6.0, 6.8, and 8.5 (Figure 3F). Interestingly, a very low Gibbs free energy (*ΔG*) was recorded for the binding assay at pH 8.5 (−969 kcal/mol) than at pH 6.0 and 6.8 (−4.96 and -4.92 kcal/mol, respectively). On the other hand, the affinity constant (*K*_*a*_) value was highest at pH 8.5 (425 ± 80 /mM) compared to *K*_*a*_ values at pH 6 and pH 6.8 (24.6 ± 1.32 and 18.9 ± 0.6 /mM, respectively). Thus, a substantially lower Gibbs free energy (*ΔG*) and 17 to 22-fold higher affinity (*K*_*a*_) values at alkaline pH suggested that manganese binds to the riboswitch element more favorably and tightly under alkaline condition.

## Discussion

The current study unfolds three hitherto unknown aspects of bacterial physiology. Firstly, we show that manganese intoxication triggers alkalization of cytoplasm, mainly altering the nitrogen metabolism (Figure 1). Secondly, precisely decoupling the intracellular enlarged manganese pool from intrinsic alkalization, we demonstrate that the concerted actions of these two factors activate the *mntP*-, and *alx*-riboswitches maximally (Figure 2 and Figure 3, Figure S7). Thirdly, we revealed how intelligently *E. coli* calibrates its metabolism under manganese stress, thereby causing intrinsic cellular alkalization, which is critical for the riboswitch-mediated manganese homeostasis (Figure 1, Figure 3A). Interestingly, alkaline pH inhibits MntH-mediated manganese import in *Salmonella enterica* (26). Since *E. coli* and *S. enterica* MntH proteins are 94% identical, it is quite plausible that the manganese-induced alkalinity in *E. coli* would also inhibit MntH function. Interestingly, the foundational study on the *mntP*-riboswitch has been performed using a mutant strain of *E. coli* (that makes MntP non-functional) (11). Using this mutant strain, they found that 1 mM manganese is enough to cause riboswitch activation (Figure 3B). Consistent with this, our study with Δ*mntP* strain also shows riboswitch activation at 1 mM manganese. However, in addition, we also used WT strain of *E. coli* to show that 8 mM manganese is required to maximally activate the riboswitch (Figure 3B).

The glutamate-glutamine cycle plays a pivotal role in acid resistance (20-22). While GlnA solely synthesizes glutamine using glutamate, two alternative pathways exist for glutamine to glutamate production. The principle one is catalyzed by GOGAT (27, 28). The other one is GlsA/B-mediated glutamine degradation, generating glutamate and ammonia (21, 29). We have previously demonstrated that manganese toxicity inactivates GOGAT (9). Thus, glutamate production via activated GlsA/B under manganese shock (Figure 2, C, D and E) liberates ammonia raising the cellular pH (Figure 4). On the other hand, a complete shutdown of glutamine to glutamate production in manganese-fed Δ*glsA*, Δ*glsB*, and Δ*mntP*Δ*glsB* mutants might affect glutamate-mediated acid resistance system that helps to raise the pH, thereby lowering cellular pH (Figure 2B). We also tested the role of glutamate dehydrogenase (*gdhA*) that catalyzes glutamate synthesis, conjugating α-ketoglutarate, and ammonia in pH elevation. However, deleting *gdhA* from the Δ*mntP* strain barely alter the growth profile, or cellular pH, under manganese stress (Figure S8), suggesting that *gdhA* plays no role under manganese stress.

**Figure 4.**
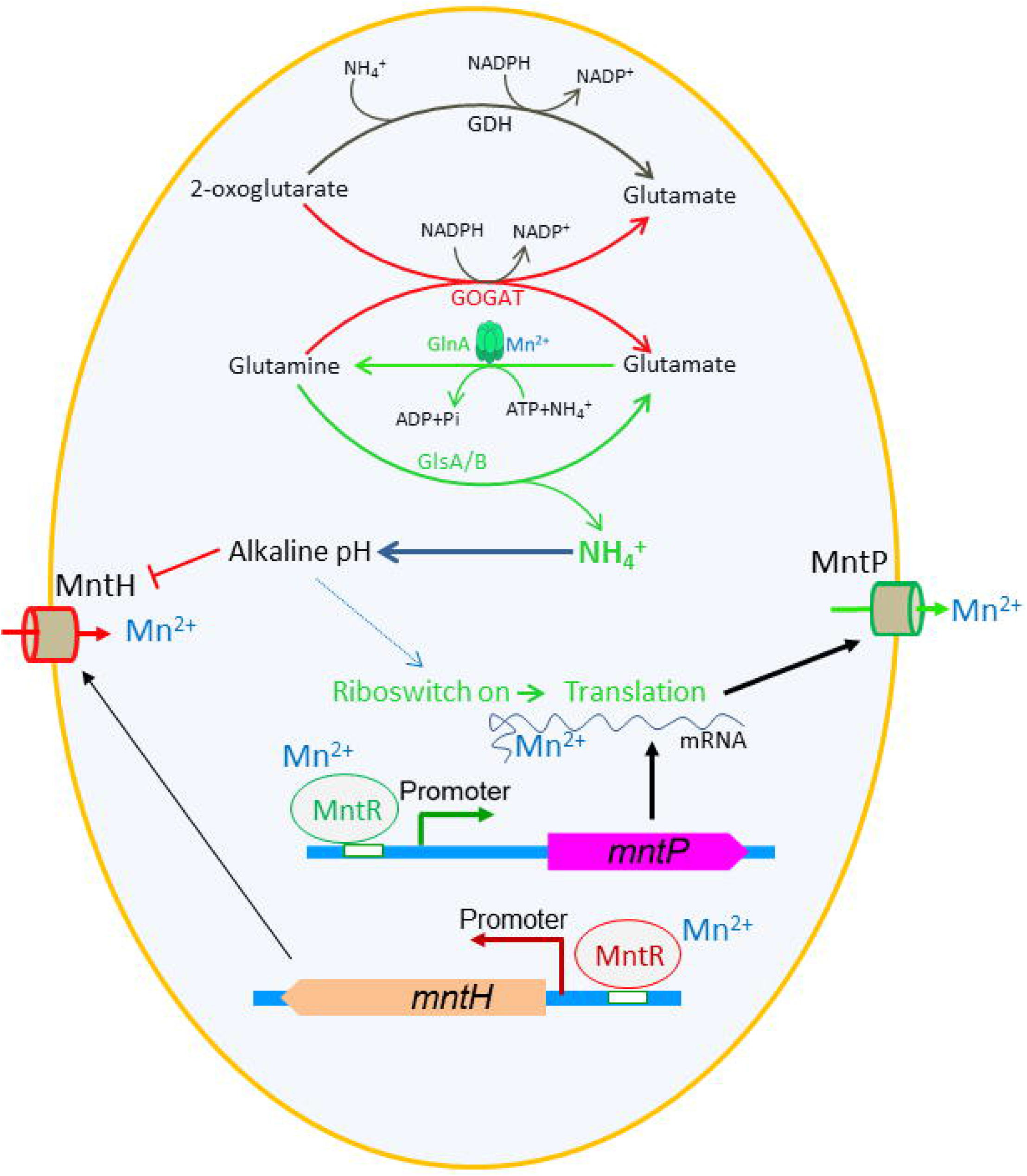
The integrated model depicts how the intrinsic cellular alkalization contributes to manganese homeostasis. Skewed functioning of the glutamate-glutamine cycle enzymes (viz. activation of GlnA and GlsA/B enzymes, and deactivation of GOGAT) under manganese stress leads to ammonia production and subsequent alkalization. The alkaline pH favors interaction between manganese and the riboswitch element to upregulate MntP expression enhancing manganese efflux. The excess manganese activates MntR to repress MntH importer. The alkaline pH also blocks MntH functioning, as reported (26), to resist manganese intoxication.

Previously, GlsA, a glutaminase that functions under acidic pH, and glutamine have been shown to constitute an acid resistance system that accelerates ammonia production to cope up with external acid shock (21, 29). However, whether ammonia production induced by GlsA action could intrinsically cause intracellular alkalization was not envisioned. Furthermore, no relevant physiological importance of GlsB, which function optimally at alkaline condition, was known so far (21, 29). The present study reveals that GlnA and GlsA/B work in concert, causing intrinsic alkalization (i.e., without extrinsic acid-shock stimulus) in *E. coli* under manganese stress. Thus, first time our study provides a rationale for the existence of two Gls proteins in *E. coli* (21, 29). Recent report suggests that deletion of glutamine synthetase (GS) in *Leishmania donovani*, affects its viability (30). *Mycobacterium tuberculosis* GS has also been considered as a potential therapeutic target to inhibit its growth (31). Whether glutamate-glutamine cycle-mediated alkalization imparts viability of these pathogens needs further investigation.

In humans, excess manganese dents the metabolic processes in basal ganglia causing manganism (32, 33). Besides, glutamine synthetase (GS) plays a pivotal role in recycling the neurotransmitter glutamate and ammonia detoxification, thereby preventing glutamate/ammonia neurotoxicity. Alteration in the expression of GS contributes to pathologies of several neurological disorders, including hepatic encephalopathy, ischemia, amyotrophic lateral sclerosis, epilepsy, Alzheimer’s and Parkinson’s diseases, traumatic brain injury, etc. (34). Glutaminases (Gls) have been reported to be involved in neuroinflammation responses of various neurological disorders (35). Thus, in neurodegenerative diseases, glutamine, ammonia, and glutamate play significant roles (36). Therefore, considering the universality of the key biochemical reactions, the elevation of pH may play critical roles in manganism and other neurological disorders.

The alkaline pH favored the interaction between manganese and the riboswitch by decreasing the *ΔG* and increasing *K*_*a*_ (Figure 3F). Corresponding dissociation constant (*K*_*d*_) values for this interaction were 41 µM, 53 µM, and 2.4 µM at pH 6.0, 6.8, and 8.5, respectively (Figure 3F). These values are well within the range of basal levels of manganese (20-30 μM) in *E. coli* (9). Especially, the *K*_*d*_ value 2.4 µM at pH 8.5 is much lower than the basal level of manganese. This observation explains how alkaline pH sensitized the riboswitch activation even in the presence of basal levels of manganese (Figure 3A). Since alkaline pH has ability to break up some Hoogstein base pairs (37), a similar scenario for *mntP*-riboswitch could facilitate further manganese binding. Especially the extended 5’-UTR portion downstream of the manganese-binding aptamer (11) could fold back to form triple helix (38) inhibiting manganese binding at lower pH. A smaller Δ*S* value at alkaline pH possibly indicates that manganese is required to make lesser disorder at alkaline pH than at acidic pH to turn the riboswitch on (Figure 3F).

The K_d_ value at pH 6.8 for *mntP*-riboswitch in our study (53 µM) matches well with the reported K_d_ values for other manganese-sensing *yybP*-*ykoY* riboswitches from *Lactococcus lactis* and *Streptococcus pneumonia* (30-40 µM and 54 ± 26 µM, respectively) (39, 40). In contrast to our finding, a previous report has shown a K_d_ value of 7 nM for manganese-sensing *mntP* riboswitch aptamer from *E. coli* at pH 6.8 (41). This low nanomolar K_d_ value is far below the intracellular concentration of manganese in WT *E. coli* (20-30 µM) (9). Therefore, this value appears to be physiologically irrelevant, as the *mntP*-riboswitch would have always been activated at that cellular level of manganese. In other words, the riboswitch activation would not have required further increase in cellular manganese level. Furthermore, where the authors have shown an enthalpy driven reaction (41), we observed an entropy driven reaction for manganese and *mntP*-riboswitch interaction (Figure 3F). The length of riboswitch sequences in our study (whole 5’-UTR), and previous report (shorter riboswitch aptamer) (41) might be the possible reason for such observed discrepancies.

## Acknowledgments

The authors are grateful to Prof Conrad Mullineaux, Queen Mary, University of London, Dr. J. Gowrishankar, CDFD, India; Dr. Rachna Chaba, IISER-Mohali, India for the generous gifts of plasmids mentioned in Table 1 and her suggestions in manuscript writing. The authors are grateful to Prof. Evgeny Nudler, NYU School of Medicine, Prof. Abhijit A. Sardesai, CDFD, India, Prof. Runa Sur, Calcutta University, India for reviewing and providing critical comments on the manuscript.

**Table 1.** The list of strains and plasmids used in this work.

| Strains and plasmids | Genotype/Features | References |
| --- | --- | --- |
| Strains |  |  |
| BW25113 | *Escherichia coli*; *rrnB3ΔlacZ4787hsdR514Δ(araBAD) 567*<br>*Δ(rhaBAD)568 rph-1* | (42) |
| $\Delta mntP$ | BW25113, $\Delta mntP::kan^R$ | (42) |
| $\Delta glnA$ | BW25113, $\Delta glnA::kan^R$ | (42) |
| $\Delta glsA$ | BW25113, $\Delta glsA::kan^R$ | (42) |
| $\Delta glsB$ | BW25113, $\Delta glsB::kan^R$ | (42) |
| $\Delta gadA$ | BW25113, $\Delta gadA::kan^R$ | (42) |
| $\Delta gadB$ | BW25113, $\Delta gadB::kan^R$ | (42) |
| $\Delta gadC$ | BW25113, $\Delta gadC::kan^R$ | (42) |
| $\Delta glsA\Delta glsB$ | BW25113, $\Delta glsA, \Delta glsB::kan^R$ | This study |
| $\Delta mntP\Delta glnA$ | BW25113, $\Delta mntP, \Delta glnA::kan^R$ | This study |
| $\Delta mntP\Delta glsA$ | BW25113, $\Delta mntP, \Delta glsA::kan^R$ | This study |
| $\Delta mntP\Delta glsB$ | BW25113, $\Delta mntP, \Delta glsB::kan^R$ | This study |
| $\Delta mntP\Delta glnA\Delta glsA$ | BW25113, $\Delta mntP, \Delta glnA, \Delta glsA::kan^R$ | This study |
| $\Delta mntP\Delta glnA\Delta glsB$ | BW25113, $\Delta mntP, \Delta glnA, \Delta glsB::kan^R$ | This study |
| $\Delta mntP\Delta gadABC$ | BW25113, $\Delta mntP, \Delta gadA, \Delta gadB, \Delta gadC::kan^R$ | This study |
| Plasmids | Features | Source |
| pET28a (+) | *kan<sup>R</sup>*; T7-promoter; IPTG inducible | Novagen |
| pET-glnA | *glnA* cloned in *NdeI* and *HindIII* sites of pET28a (+) vector | This study |
| pCL1920 | *amp<sup>R</sup>*; Lac-promoter, IPTG inducible | A gift from Prof. J. Gowrishankar (43) |
| pAH125 | *kan<sup>R</sup>*; LacZ reporter | A gift from Prof. R. Chaba (44) |
| pINT | *amp<sup>R</sup>*; λ-integrase | A gift from Prof. R. Chaba (44) |
| pBAD-GFPmut3* | *amp<sup>R</sup>*; Lac-promoter, arabinose inducible | Prof C. Mullineaux (23) |
| pGlnA | pCL1920 with *glnA* at *NdeI* and *HindIII* | This study |
| pBAD/*Myc*-His A | *amp<sup>R</sup>*; cloning vector | Invitrogen |
| pDAK1 | pBAD/*Myc*-His A; Two *NdeI* sites were mutated and *NcoI* site was replaced by *NdeI* | This lab |
| pGlsA | pDDAK1 with *glsA* at *NdeI* and *HindIII* | This study |
| pGlsB | pDDAK1 with *glsB* at *NdeI* and *HindIII* | This study |
| NOTE: *kan<sup>R</sup>*, kanamycin resistance; *amp<sup>R</sup>*, ampicillin resistance and *cm<sup>R</sup>*, chloramphenicol resistance. |  |  |

## Author contributions

Conceptualization: D.D., Methodology: A.K., R.K.M., V.K, A.A. and D.D. Investigation: A.K., R.K.M., V.K, A.A. and D.D. Visualization: A.K. and D.D. Funding acquisition: D.D. Project administration: D.D. Supervision: D.D. and A.A. Writing – original draft: A.K. Writing – review & editing: A.K. A.A. and D.D.

## Data availability

The microarray data generated and used in this study have been deposited in Gene expression Omnibus (GEO) database with accession number GSE186657.

## Materials and Methods

### Bacterial strains, plasmids, media, and growth conditions

The wild-type BW25113 (WT) and the KEIO knockout mutant strains of *E. coli* used in this study are listed (Table 1). The knockouts were verified by PCR, freshly transduced into the WT background by P1 phage, and sequenced to confirm the deletion. The double and triple mutants were generated by P1 phage transduction. P_*glsA*_-*lacZ* transcriptional fusion construct was made by fusing the entire *glsA* promoter region beginning with 220 nucleotides upstream of the transcription site and the 68 nucleotides 5′-UTR of *glsA* to *lacZ*. Similarly, we constructed two more translational fusion reporters, P_T7A1_-5′-UTR_*mntP-*17-codons_ –*lacZ* and P_T7A1_-5′-UTR_*alx-*17-codons_ – *lacZ*. For this, the entire 5′-UTR and first 17 codons of the *mntP* or *alx* genes were fused with *lacZ*. Further fusion PCRs were performed to attach the T7A1 promoter upstream of these reporters. We also made P_T7A1_-*lacZ* reporter as a negative control to check the effects of different pH on T7A1 activity. The reporter constructs were cloned into pAH125 vector (44). The cloned constructs were then integrated into the genome of WT strain using pINT helper plasmid and screened for single integrant, as described (44). The genotype was transduced to various desired strains of *E. coli* by P1 transduction. The gifted and constructed plasmids in this study are mentioned in Table 1. The oligonucleotides used in the study are mentioned in Table S1. All studies were done in LB broth and LB-agar media at 37°C.

### Growth curve analysis

The primary culture was grown overnight at 37°C with shaking. For growth analysis, the cultures were diluted to 0.6 OD_600_, and 1% of the diluted culture was inoculated in 1 ml of LB broth in the presence and absence of desired concentrations of manganese. Growth curve analysis was done using a Bioscreen C growth analyzer (Oy growth curves Ab Ltd).

### Viability assay

To check the survivability under extreme manganese stress, 1% overnight cultures were inoculated in LB media supplemented with 10 mM manganese. After incubating for indicated times, the cultures were diluted as required and plated on the LB agar surface. The number of colonies were counted and compared with the initial number of cells determined from the untreated counterparts.

### Microarray experiments

The saturated overnight cultures of *E. coli* strains were inoculated in the fresh LB medium at 1:100 dilution and grown initially for 1 to 1.5 hours to get the O.D. about 0.3 and then grown again for 2.5 hours in the presence or absence of 1 mM manganese at 37 °C before harvesting the cell pellets. The pellets were washed with normal saline (0.9%) and stored by dissolving in RLT buffer (Sigma) before further processing. The microarray was done on Agilent based customized platform from Genotypic Technology, Bangalore. On average, three probes were designed for each gene. RNA extraction, quality control, microarray labelling, hybridization, scanning, and Data analyses were done, as mentioned previously (44). The microarray data generated and used in this study have been deposited in Gene expression Omnibus (GEO) database with accession number GSE186657.

### Estimation of intracellular glutamine and glutamate levels

WT and Δ*mntP* cells were grown in the presence or absence of 1 mM manganese. The cell pellets were washed with PBS and resuspended in 50% cold methanol. Subsequently, cells were subjected to three freeze-thaw cycles in liquid nitrogen followed by sonication. The supernatants were collected and dried using a SpeedVac vacuum concentrator. LC-MS/MS analyses and quantification was done at metabolomics facility, National Institute of Plant Genome Research, New Delhi, India, as per protocol (45). The amino acid levels were expressed as nmol/mg of wet cell pellet weights.

### Estimation of intracellular pH

The intracellular pH was estimated, as described (24). Briefly, 1% of overnight primary cultures of the desired strains of *E. coli* harboring pAra-TorA-GFPmut3*plasmid (23, 24) were inoculated in 200 ml of fresh LB broth to achieve an OD_600_ 0.125. The cells were grown in the presence or absence of supplemented manganese or glutamine (at desired concentrations indicated in the result section) for 2 hours, and then GFP was induced with 0.003% L-arabinose. 0.2 mM IPTG was used to induce GlnA from pGlnA-complemented strains. The cells were grown for another 2 hours at 37 □C after induction. The individual bacterial cell pellets were collected and washed with 1X PBS. The cell pellets were divided into 7 equal portions. 6 portions of pellets were permeabilized by 20 mM sodium benzoate and incubated with 6 different buffers (pHs were 5.0, 6.0, 7.0, 7.5, 8.0, and 8.5) for 10 minutes, so that the intracellular pH was equilibrated with applied external pH. GFP excitation was measured from 480 to 510 nm, using an emission wavelength of 545 nm, and the standard curves were generated, as described (24). The remaining cell pellet was dissolved in PBS to measure the GFP fluorescence directly. The intracellular pH was determined from the standard curve equations.

### β-galactosidase assay

For the β-galactosidase assay, 1% of the overnight culture of bacterial strains were inoculated in fresh LB broth (initial OD_600_ was 0.125), and grown in the presence or absence of 1 mM manganese for 4 hours at 37 □C. The cell pellets were washed two times with Z-buffer (60 mM Na_2_HPO_4_, 40 mM NaH_2_PO_4_, 10 mM KCl, and 1 mM MgSO_4_), and the β-galactosidase assay was performed as described (46). Background β-galactosidase activity from the reporter-less isogenic *E. coli* strains was determined and subtracted.

For the β-galactosidase assay under exogenous alkaline and acid shocks were performed, as described below. *E. coli* was grown in LB broth for 3 hours at 37 □C and then subjected to alkaline or acid shock. For this, the culture pH was adjusted to either 9.5 (using 4M NaOH) or 4.0 (using 1M HCl) and grown for 1 hour at 37 □C. In one set of assays, we measured the intracellular pH. In another set of assays, we performed β-galactosidase assays to know the riboswitch activity. The riboswitch activities were also checked by growing the *E. coli* cells in different buffered (pH 5.0, 7.0 and 8.5) LB broth.

### Western blotting experiments

Untreated and manganese-treated cell pellets were harvested, washed with 1X PBS, and then lysed with B-PER® bacterial protein extraction reagent (Thermo Scientific) followed by sonication. The total protein level was estimated by using the Bradford assay kit (Bio-Rad). 30 µg of total cellular proteins from the individual samples were subjected to SDS-PAGE. The proteins were transferred to nitrocellulose membrane and stained with Ponceau S to visualize the resolution and equal loading in the PAGE. Western blotting was performed using polyclonal rabbit primary antibodies raised against purified GlnA protein, and HRP conjugated secondary goat anti-rabbit polyclonal antibodies (Sigma Aldrich). The blots were developed by Immobilon® Forte Western HRP substrate (Millipore).

### RNA synthesis by in vitro transcription

The template DNA for in vitro transcription was prepared by fusing the T7 promoter and the *mntP* riboswitch element, including 357 bases. The RNA was synthesized using the T7 RiboMax express large-scale RNA production system, Promega. To remove the DNA template, the RQ1 RNase-free DNase was added to the reaction mixture to a concentration of 1 unit per microgram of template DNA and incubated for 15 minutes at 37 □C. The reaction mixture was then extracted with phenol-chloroform and ethanol precipitated. The eluted RNA was then passed through the P-6 column to remove the unincorporated nucleotides.

### Isothermal titration calorimetry

A MicroCal Auto-iTC200 calorimeter, MicroCal Inc., was used for calorimetric measurements to probe the interaction of *mntP* riboswitch element with manganese in different pH. Prior to the ITC experiments, the *mntP* riboswitch RNA was equilibrated in the ITC buffers [30 mM HEPES (for pH 6 and 6.8) or 30 mM Tris-HCl (for pH 8.5), 150 mM NaCl] and then heated to 95 °C for 2 min and placed on ice for 10 min. 5 mM MgCl_2_ was then added to the RNA solutions and incubated at 21 °C for 15 min. The RNA samples were centrifuged for 20 mins at 13000 rpm. MnCl_2_ was dissolved in ITC buffers and filtered. Experiments were performed with 4 μM RNA (cell) and 75 μM of MnCl_2_ (syringe) at 25 °C. MnCl_2_ was also injected into the respective ITC buffers to get background enthalpy changes and subtracted from the experimental binding data. Data were fitted using a one-site binding model fixing stoichiometry (N) at 2 using MicroCal Origin software. Individual ITC experiments were performed 3 times.

### Ammonia liberation assay

The ammonia liberation from *E. coli* strains was estimated using the ab83360 Ammonia Assay Kit (Abcam). As mentioned previously, the cells were grown before harvesting the pellets. Pellets were washed with cold 1X PBS. 10 mg of cells were resuspended in 100 µl of assay buffer. The samples were then centrifuged at 13000 rpm for 5 minutes at 4 □C. The supernatant was collected in a clean tube and kept on ice. 50 µl of reaction mix was added to standards and the sample wells in a 96 well plate and incubated in the dark at 37°C for 60 minutes. The plate reading was immediately taken at OD_570_ nm in a microplate reader.

### Glutaminase A activity assay

To assess the activity of GlsA enzyme, different *E. coli* strains were grown in the presence and absence of manganese. The cells were then pelleted and washed with 1X PBS. The pellets were resuspended in 1X PBS by normalizing according to their respective weight. 300 μl of cell suspension (approximately 30 mg of wet cell weight) was further taken from each sample and centrifuged. The pellets were dissolved in 200 μl of GlsA assay solution containing 1 g/l L-glutamine, 0.25 g/l bromocresol green, 90 g/l NaCl, and 3 ml/l Triton X-100. pH was adjusted to 3.2 with HCl. The resuspended cell pellets were mixed thoroughly and incubated at 37 □C for 1, 3, 5, and 10 minutes. We recorded the color changes. The absorbance from 400 nm to 700 nm was recorded after collecting the supernatant from each time-point using a Synergy H1 Hybrid plater reader, BioTek.

To generate a correlation between 620/420 ratios of bromocresol green dye and pH, NaOH was gradually added to the GlsA assay solution and pH elevation was recorded after each addition. A portion of the solution was also taken out at after each addition spectral scan from 400-700 nm wavelengths. The spectrum values were normalized by considering OD values 0.1 at 515 nm, the isosbestic point (515 nm) of bromocresol green dye.

### ICP-MS analyses to determine cellular manganese levels

The cells were grown in the presence and absence of manganese for 4 hours at 37 °C and then pelleted. The pellets were washed twice with 1X PBS supplemented with 1 mM EDTA. The final washing was done with 1X PBS. The washed cells pellets were digested by adding 500 µl of 30% H_2_O_2_ and 3.5 ml of concentrated HNO_3_. The samples were heated over flame for 3 minutes and made up the final volume of 10 ml with MilliQ water. Digested samples were centrifuged at 10000 rpm for 20 minutes. The manganese levels in the supernatants were determined using ICP-MS at Punjab Biotechnology Incubator, Mohali, India. The cellular metal concentrations were calculated by considering a total cellular protein concentration of 300 mg/ml, as described (47).

### Statistical analyses

The experiments were performed 3-6 times. Data analyses, normalization, etc. were done using Microsoft Excel, Microcal Origin and GraphPad Prism software. Densitometry of the western blots was done using Image J software. The graphs were plotted as mean ± S.D. The P values were determined from the unpaired t-test.

