## Supplementary material for "An intrinsic alkalization circuit turns on *mntP*-riboswitch under manganese stress in *Escherichia coli*": Supplemetary Materials

**This PDF file includes:**

Figs. S1 to S8

Tables S1

**Figure S1. The growth analyses of different *E. coli* strains in the presence or absence of manganese stress. A.** The growth profile of *E. coli* WT, Δ*mntP*, Δ*glnA*, and Δ*mntP*Δ*glnA* strains in the absence of supplemental manganese. **B-E.** The growth profiles of the same *E. coli* strains under 0.25 mM (**B**), 0.5 mM (**C**), 0.75 mM (**D**), and 1 mM (**E**) supplemental manganese.

**Figure S2. Effect of manganese on the growth of *E. coli* strains. A.** The comparative growth profiles of WT and Δ*mntP* strains revealed that 8mM $MnCl_2$ (Mn) has the same effect on growth of WT cells as 1mM manganese on the growth of Δ*mntP* cells. **B**. The panel represents that while 0.5 mM manganese mildly inhibits growth of Δ*mntP* cells, it severely affects the growth of Δ*mntP*Δ*glnA* strain. The multicopy expression of *glnA* from pGlnA plasmid (*glnA* cloned in pCL1920 vector) rescued the growth defects of the double mutant. **C.** The panel shows that the survivability ofΔ*mntP* and Δ*mntP*Δ*glnA* strains under 10 mM manganese.

**Figure S3. Growth defects of Δ*glsA or* Δ*glsB* containing double and triple mutants on the LB-agar plate. A.** Control plates showing that the designated single and double mutants grow normally on the LB-agar surface. **B.** The plates showing the compromised growth of Δ*glnA*Δ*glsA*, Δ*mntP*Δ*glsA*, and Δ*mntP*Δ*glnA*Δ*glsA* mutants at 12 and 18 hours of incubation. **C.** The plates showing the compromised growth of Δ*glnA*Δ*glsB*, and Δ*mntP*Δ*glnA*Δ*glsB* mutants at 12 and 18 hours of incubation. **D.** The multicopy expression of *glsA* or *glsB* in the Δ*glnA*Δ*glsA*, and Δ*glnA*Δ*glsB* strains, respectively, rescued the growth defects.

**Figure S4. Kinetics of color change of bromocresol green indicator dye with pH. A.** Color of the indicator dye at different pH has been shown. **B**. Absorption spectra of the indicator dye at different pH values show that the absorption at 620 nm increases, while the absorption at 420 nm decreases with increasing pH. **C.** Ratio of the two absorption peaks (620/420) were plotted at different pH.

**Figure S5. GlsA activity in the untreated and the manganese-treated *E. coli* strains. A.** Kinetics of GlsA-catalyzed ammonia base liberation from the *E. coli* strains were monitored at 1, 3, 5, 10 minutes time-points using bromocresol green dye. **B. i.** The panel is identical to the Figure 2E showing the increasing 620/420 ratios with time for the various *E. coli* strains grown, as indicated. **ii.** The approximate pH at different time-points were calculated from the 620/420 ratios (using fig S3C data) and plotted.

**Figure S6. The riboswitch activity under different buffered LB broth. A.** The *ΔmntP* mutant was grown in different buffered LB broth with varying pH (5.0, 7.0 and 8.5) and in the presence or absence of manganese to show that the alkaline pH can activate the *mntP*-riboswitch independent of manganese stress. **B.** $P_{T7A1}$-lacZ reporter construct show that the promoter activity was not influenced by the pH variations.

**Figure S7. Activation of *alx*-riboswitch under manganese stress and alkaline shock.** The *ΔmntP* mutant was grown in different buffered LB broth with varying pH (5.0, 7.0 and 8.5) and in the presence or absence of manganese to show that the alkaline pH can activate the *alx*-riboswitch independent of manganese stress.

**Figure S8. GdhA plays no significant role under manganese stress. A.** The growth curves reveal that Δ*mntP* and Δ*mntPΔgdhA* strains grows similarly under 0.5 mM manganese stress. **B.** 1mM manganese shock elevated the intracellular pH from 7.5 to 8.3 in the Δ*mntPΔgdhA* mutant, which was similar to the pH in the unfed and 1 mM manganese-fed Δ*mntP* strain (Figure 1E).

### Table S1. The list of oligonucleotides used in this study

| Oligonucleotides | | |
|---|---|---|
| Oligo Names | Sequence | Details |
| AK_glnA_F_28a | GCTACTTCATATGTCCGCTGAACAC GTAC | *glnA* forward primer with *NdeI* site. To clone in pET28a (+) vector |
| AK_glnA_R_28a | CATGCTAAGCTTTTAGACGCTGTAG TACAGCTC | *glnA* everse primer with *HindIII* site. To clone in pET28a (+) vector |
| AK_glnAF_1920 | GCTACTTAAGCTTGATGTCCGCTGA ACACGTAC | *glnA* gene Forward primer with *HindIII* site. To clone in pCL1920 vector |
| AK_glnAR_1920 | CATGCTGGTACCTTAGACGCTGTAG TACAGCTC | *glnA* gene Reverse primer with *KpnI* site. To clone in pCL1920 vector |
| T7A1_F | CACTGAGGTACCTCCAGATCCCGAA AATTTATCAA | T7A1 promoter Forward primer With *KpnI* site. Used in riboswitch fusion construct preparation |
| T7A1_R | TCGAGAGGGAGCAACCGCTGGAGTT C | T7A1 promoter Reverse primer used in riboswitch fusion construct preparation |
| AK_1 mntP_T7A1 overhang | AGCAACCGCTGGAGTTCGTTAAGGT TGCGCCCTC | *mntP* riboswitch Forward primer with T7A1-reverse primer overhang. Used in riboswitch fusion construct preparation. |
| AK_2 mntP_rev | GACAATGTCCTGACCGG | *mntP* riboswitch Reverse primer. Used in riboswitch fusion construct preparation. |
| AK_3 lacZ_com_F | TTATGGATACCCCGGTCAGGACATT GTCATGACCATGATTACGGATT | *lacZ* gene Forward primer with *mntP* riboswitch reverse primer overhang. Used in riboswitch fusion construct preparation. |
| AK_4 lacZ _R | GCATGGCTAGCTTCTTCGTCTGTTTC TACTGGTATTGGC | *lacZ* gene Reverse primer with *NheI* site. Used in riboswitch fusion construct preparation. |
| AK_glsAlcF | GCATCAGGTACCCAGGGTCAGGTCG ATAG | *glsA* gene Forward primer with *KpnI* site. Used in making lacZ reporter promoter fusion. |
| AK_glsAlcR | GCTACGTCGACCTTTTGTTAACTCC | *glsA* gene Reverse primer with *HindIII* site. Used in making lacZ reporter promoter fusion. |
| AK_T7mntP_F | GAGATCTCGATCCCGCGAAATTAAT ACGACTCACTATAGGGGGTTAAGGT TGCGCCCTC | T7 promoter primer with *mntP* riboswitch overhang for in vitro transcription template preparation. |
| AK17 | GAATTCCTAGGTTCGCAATGCTTCA G | *mntP* riboswitch reverse primer for in vitro transcription template preparation. |
| SB3 | GCATGGCATATGTTAGATGCAAACA AATTACAGC | *glsA* gene Forward primer with *NdeI* site. To clone in pDAK1 vector. |
| SB4 | GCTTACAAGCTTCAGCCCTTAAACA CG | *glsA* gene Reverse primer with *HindIII* site. To clone in pDDAK1 vector |

| | | |
|---|---|---|
| SB5 | GCATGGCATATGGCAGTCGCCATGG ATAATGC | *glsB* gene Forward primer with *NdeI* site. To clone in pDAK1 vector. |
| SB6 | GCTTACAAGCTTCGAGAGACTGCAT TAATAAACC | *glsB* gene Reverse primer with *HindIII* site. To clone in pDAK1 vector |

**Supplementary Figures**

Figure S1

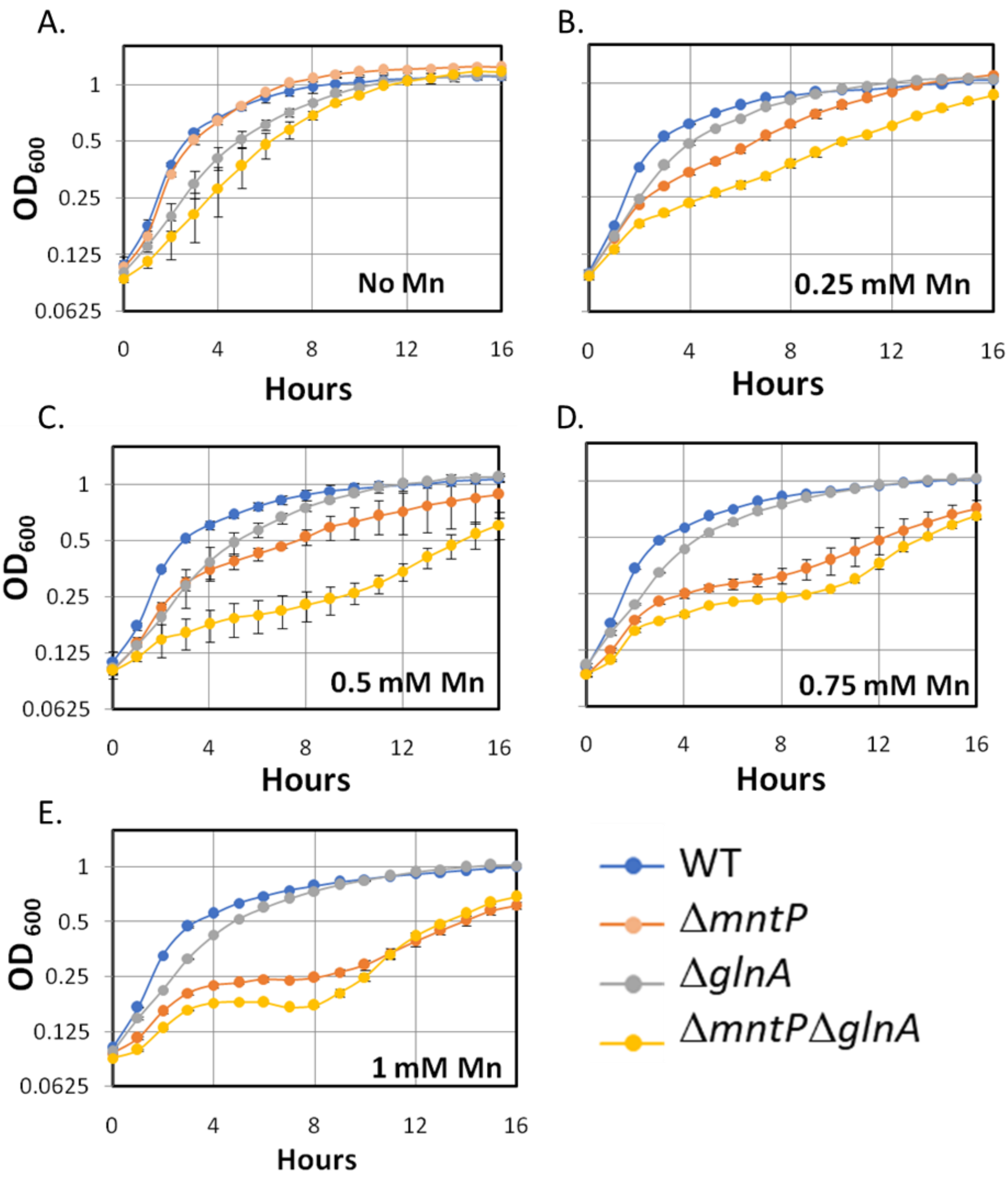

**Figure S2**

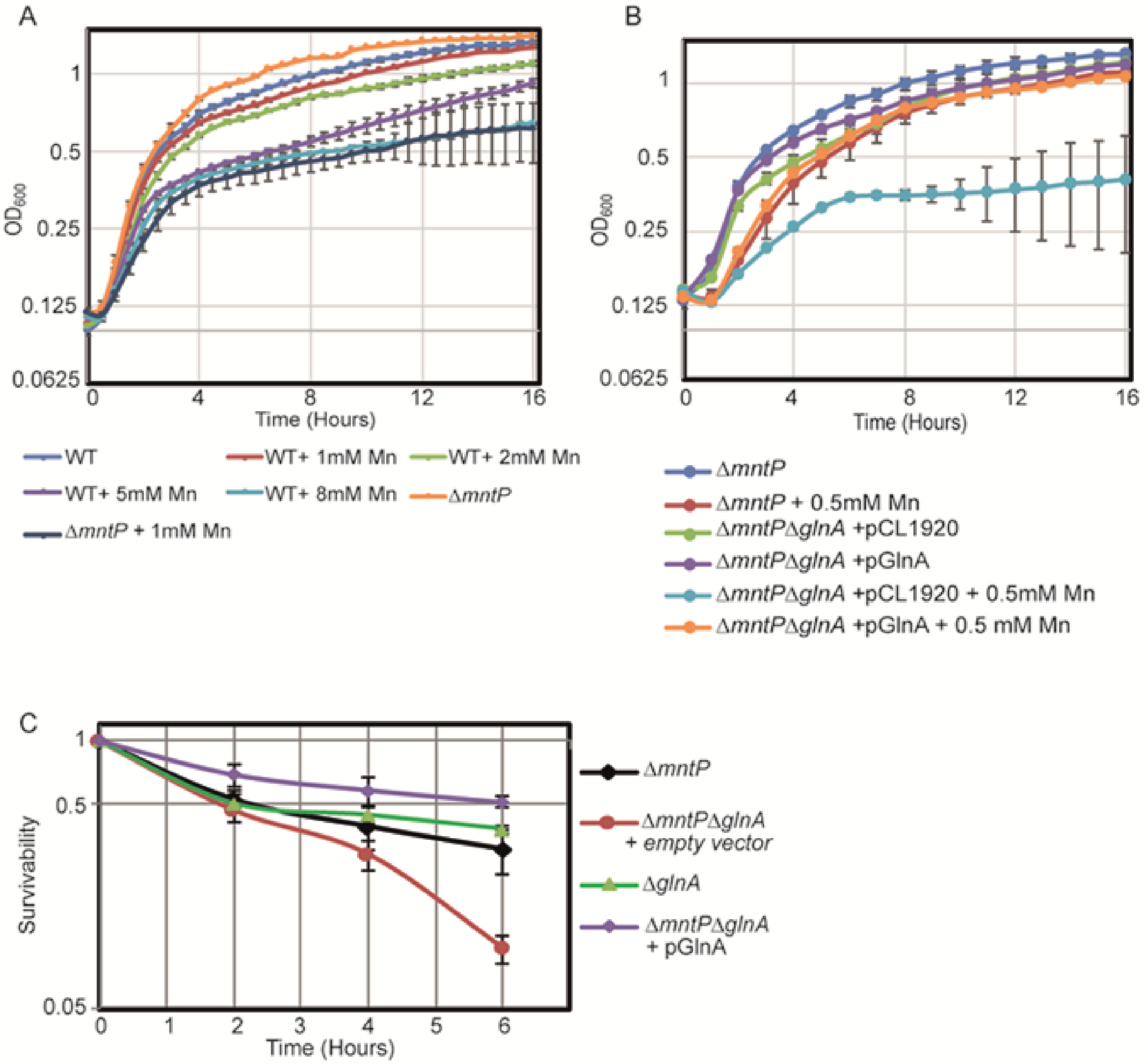
A
OD600
1
0.5
0.25
0.125
0.0625
0
4
8
12
16
Time (Hours)
WT
WT+ 1mM Mn
WT+ 2mM Mn
WT+ 5mM Mn
WT+ 8mM Mn
ΔmntP
ΔmntP + 1mM Mn
B
OD600
1
0.5
0.25
0.125
0.0625
0
4
8
12
16
Time (Hours)
ΔmntP
ΔmntP + 0.5mM Mn
ΔmntPΔglnA +pCL1920
ΔmntPΔglnA +pGlnA
ΔmntPΔglnA +pCL1920 + 0.5mM Mn
ΔmntPΔglnA +pGlnA + 0.5 mM Mn
C
Survivability
1
0.5
0.05
0
1
2
3
4
5
6
Time (Hours)
ΔmntP
ΔmntPΔglnA
+ empty vector
ΔglnA
ΔmntPΔglnA
+ pGlnA

**Figure S3**

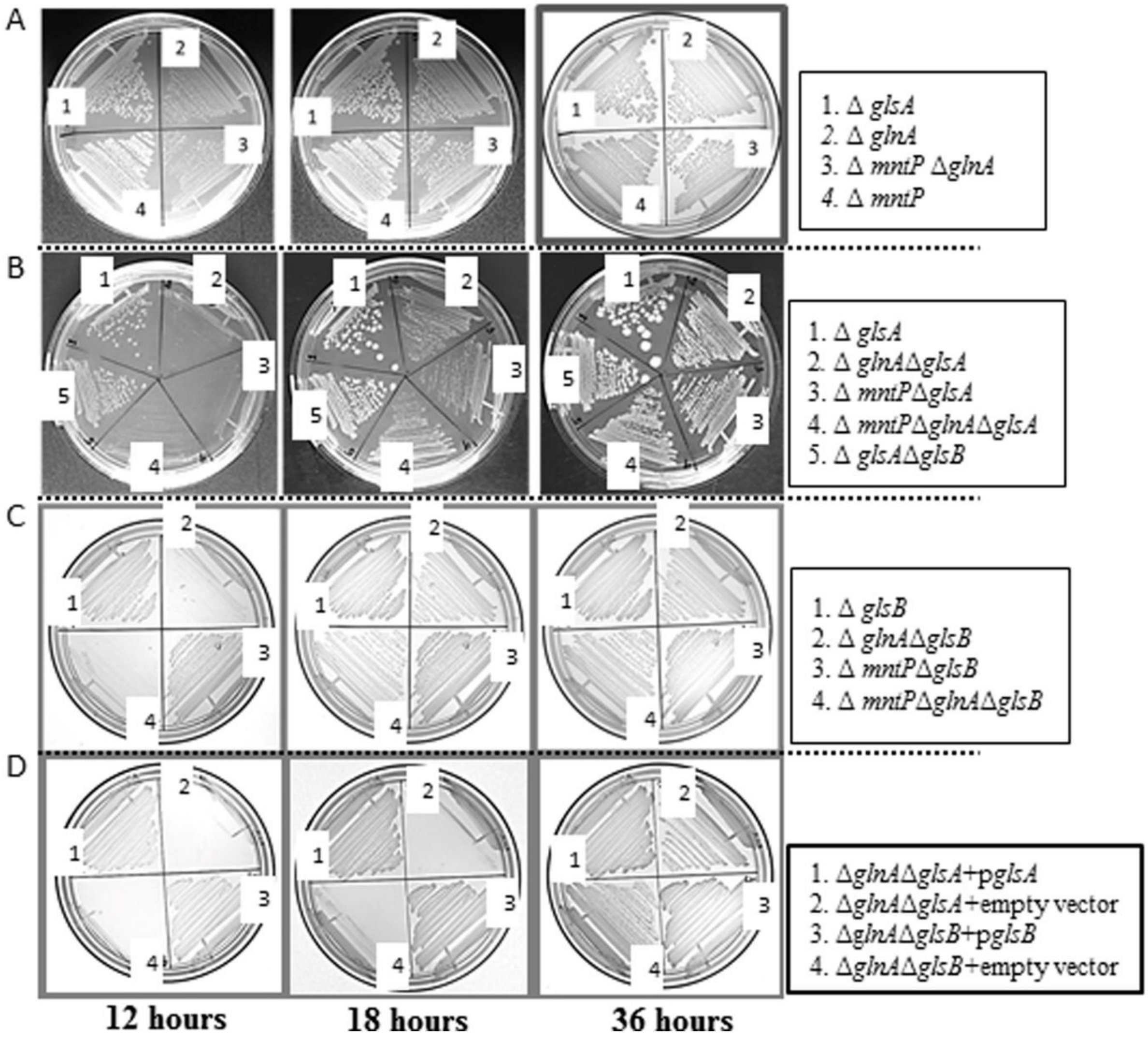
A
1. Δ glsA
2. Δ glnA
3. Δ mntP ΔglnA
4. Δ mntP
B
1. Δ glsA
2. Δ glnAΔglsA
3. Δ mntPΔglsA
4. Δ mntPΔglnAΔglsA
5. Δ glsAΔglsB
C
1. Δ glsB
2. Δ glnAΔglsB
3. Δ mntPΔglsB
4. Δ mntPΔglnAΔglsB
D
1. ΔglnAΔglsA+pglsA
2. ΔglnAΔglsA+empty vector
3. ΔglnAΔglsB+pglsB
4. ΔglnAΔglsB+empty vector
12 hours
18 hours
36 hours

**Figure S4**

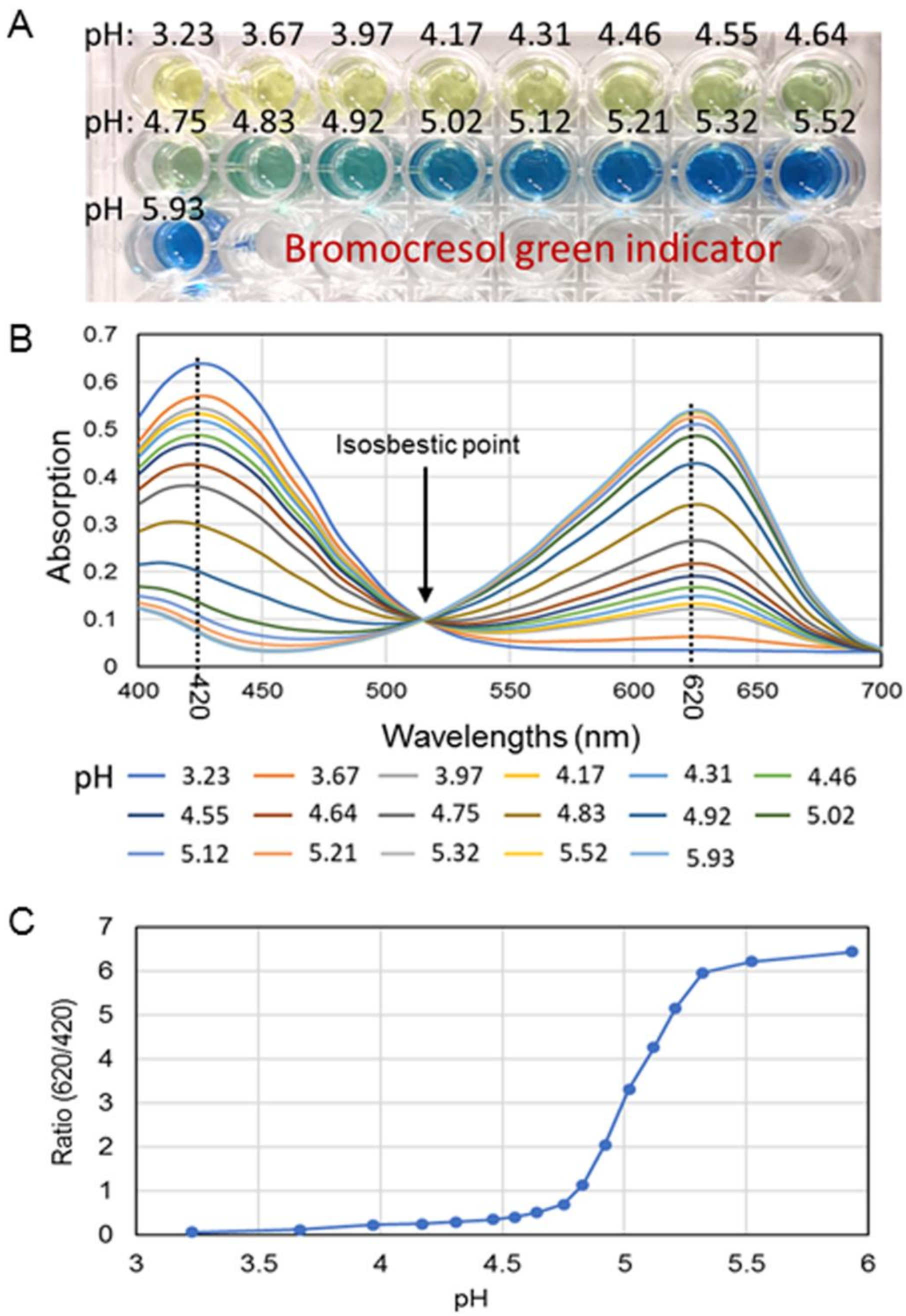

A
pH: 3.23 3.67 3.97 4.17 4.31 4.46 4.55 4.64
pH: 4.75 4.83 4.92 5.02 5.12 5.21 5.32 5.52
pH 5.93
Bromocresol green indicator
B
Absorption
Isosbestic point
Wavelengths (nm)
420
620
pH 3.23 3.67 3.97 4.17 4.31 4.46 4.55 4.64 4.75 4.83 4.92 5.02 5.12 5.21 5.32 5.52 5.93
C
Ratio (620/420)
pH

### Figure S5

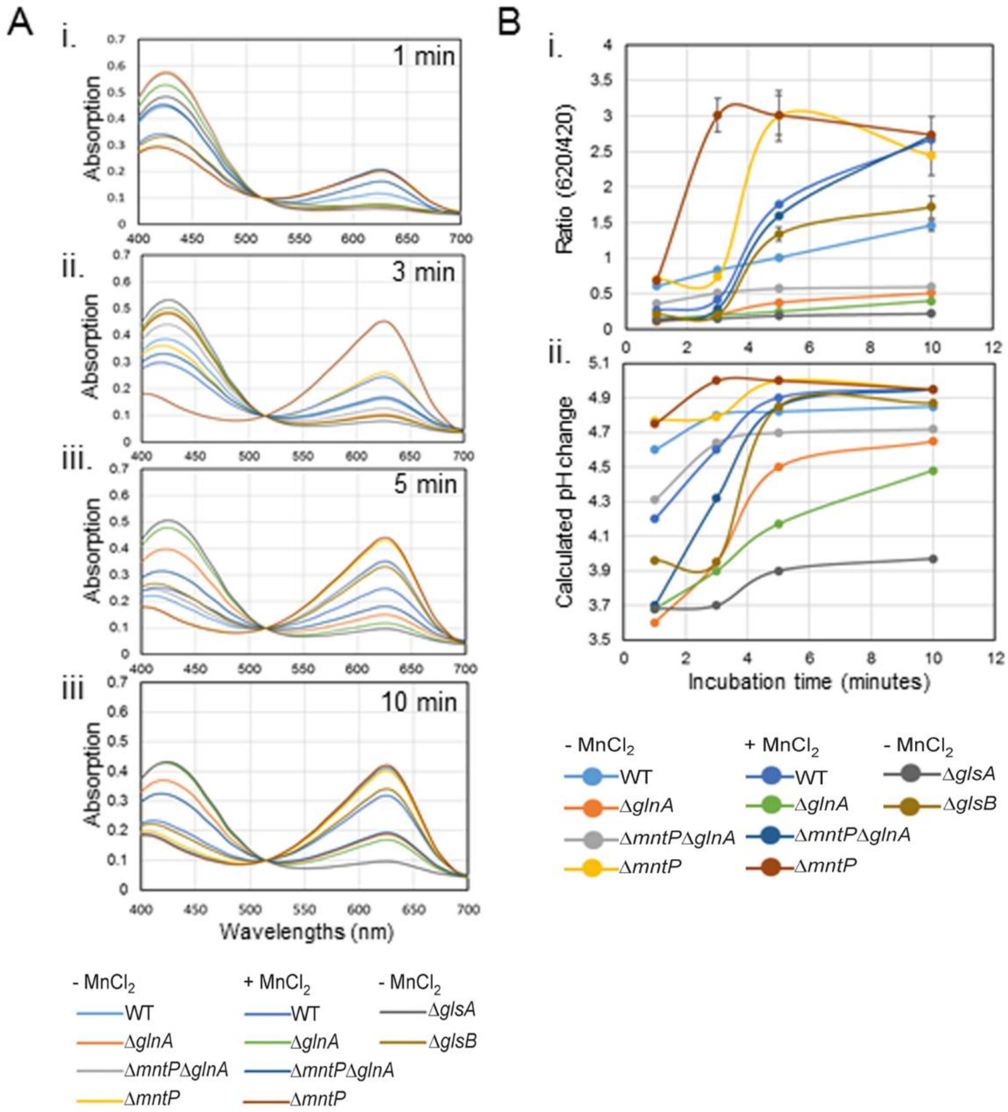
A
i.
1 min
ii.
3 min
iii.
5 min
iii
10 min
Absorption
Wavelengths (nm)
B
i.
Ratio (620/420)
ii.
Calculated pH change
Incubation time (minutes)
- MnCl2
+ MnCl2
- MnCl2
WT
ΔglnA
ΔmntPΔglnA
ΔmntP
WT
ΔglnA
ΔmntPΔglnA
ΔmntP
ΔglsA
ΔglsB

**Figure S6**

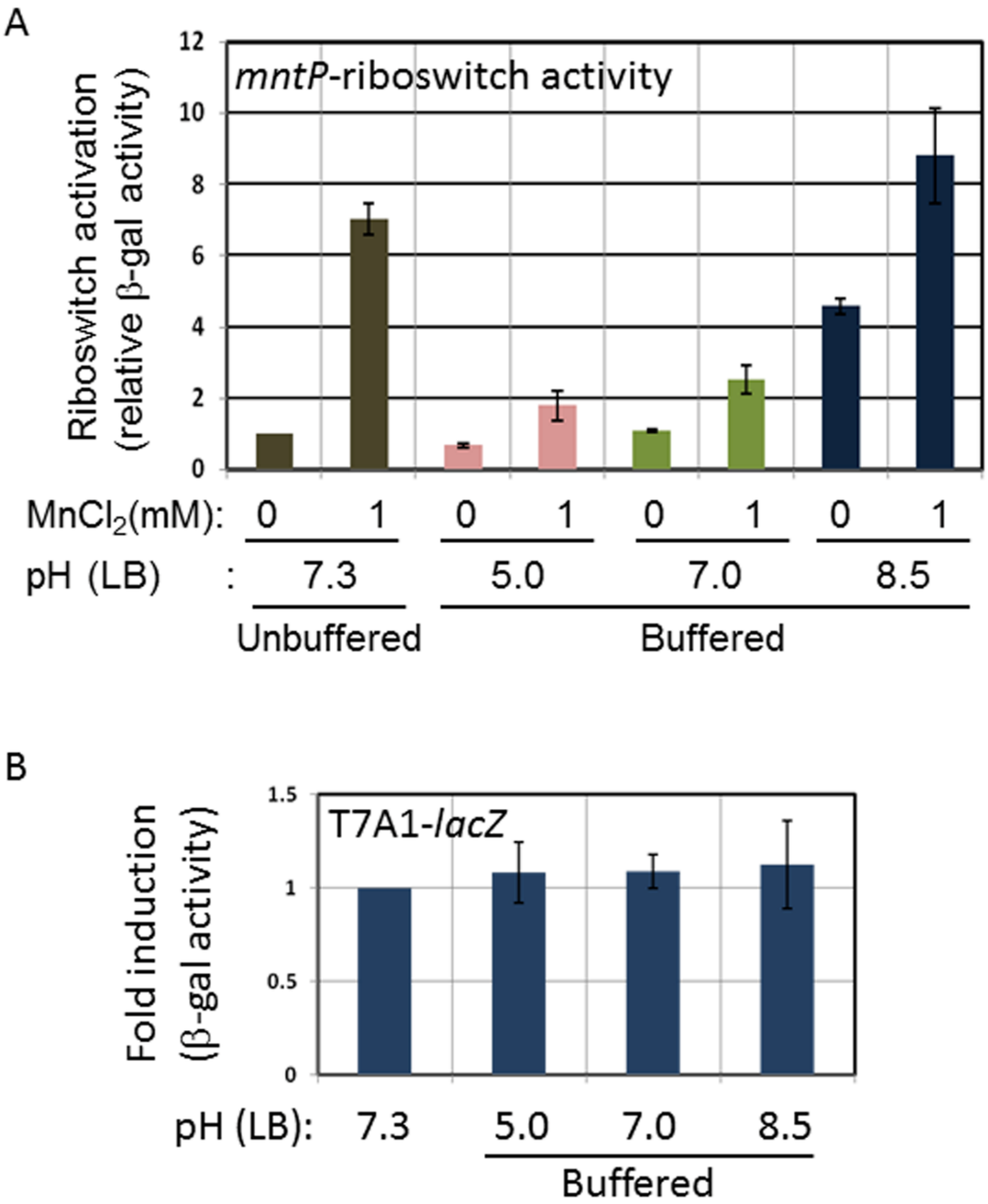
A
mntP-riboswitch activity
Riboswitch activation
(relative β-gal activity)
0
2
4
6
8
10
12
MnCl2(mM): 0 1 0 1 0 1 0 1
pH (LB) : 7.3 5.0 7.0 8.5
Unbuffered
Buffered
B
T7A1-lacZ
Fold induction
(β-gal activity)
0
0.5
1
1.5
pH (LB): 7.3 5.0 7.0 8.5
Buffered

**Figure S7**

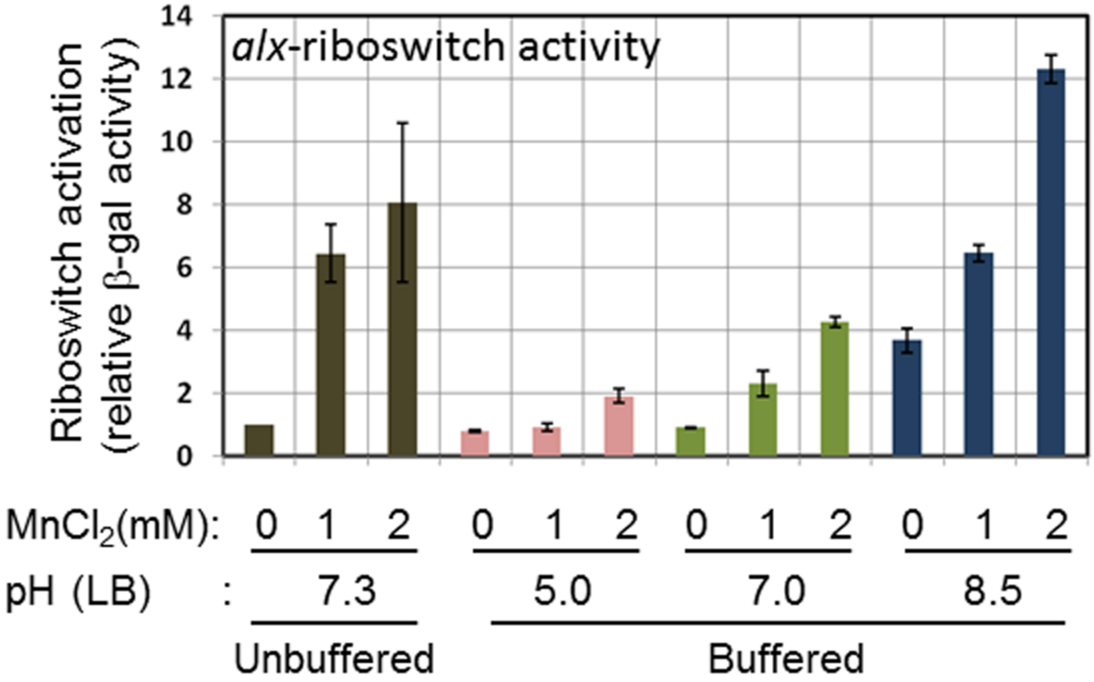
alx-riboswitch activity
Riboswitch activation
(relative β-gal activity)
14
12
10
8
6
4
2
0
MnCl2(mM): 0 1 2 0 1 2 0 1 2 0 1 2
pH (LB) : 7.3 5.0 7.0 8.5
Unbuffered
Buffered

**Figure S8**

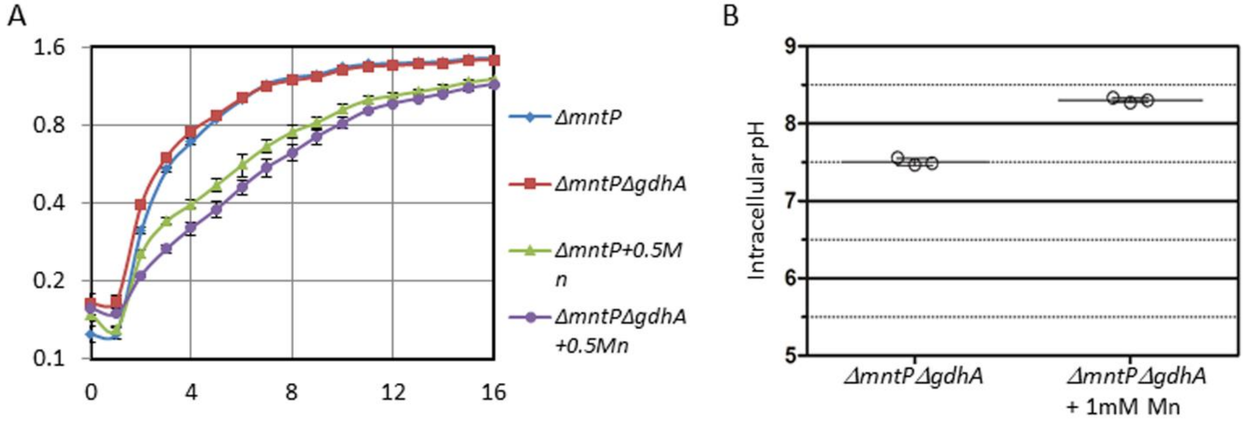
A
1.6
0.8
0.4
0.2
0.1
0
4
8
12
16
ΔmntP
ΔmntPΔgdhA
ΔmntP+0.5Mn
ΔmntPΔgdhA +0.5Mn
B
9
8
7
6
5
Intracellular pH
ΔmntPΔgdhA
ΔmntPΔgdhA + 1mM Mn
